# The landscape and diagnostic potential of T and B cell repertoire in Immunoglobulin A Nephropathy

**DOI:** 10.1101/316216

**Authors:** Chen Huang, Xuemei Li, Jinghua Wu, Wei Zhang, Shiren Sun, Liya Lin, Xie Wang, Hongmei Li, Xiaolei Wu, Peng Zhang, Guoshuang Xu, Hanmin Wan, Hongbao Liu, Yuzhen Liu, Dapeng Chen, Li Zhuo, Huanming Yang, Jian Wang, Ling Wang, Xiao Liu

**Affiliations:** Department of Nephrology, Xijing Hospital, The Fourth Military Medical University, Xi’an, China; BGI-Shenzhen, Shenzhen, China; BGI-Tianjin, Tianjin, China; Department of Nephrology, China-Japan Friendship Hospital, Beijing, China; Department of vascular and endocrine surgery, Xijing hospital, Forth military medical university, Xi’an, China

## Abstract

Immunoglobulin A Nephropathy (IgAN) is the most common glomerulonephritis worldwide. In IgAN, immune complex deposite in glomerular mesangium, which induce inflammation and affect the kidney’s normal functions. However, the exact pathogenesis of IgAN is still incompletely understood. Further, in current practice the clinical diagnosis relies on needle biopsy on renal tissue. Therefore, a non-invasive method for clinical diagnosis and prognosis surveillance of the disease is in high demand. In this paper, we investigated both the T cell receptor bata chain (TCRB) and immunoglobulin heavy chain (IGH) repertoire of kidney infiltrating and circulating lymphocytes of IgAN patients by immune repertoire high throughput sequencing. We found that the features of TCRB and IGH in the renal tissues were remarkably different from that in blood, including a decreased repertoire diversity and increased IgA and IgG frequency, and more activated B cells. The CDR3 length of PBMC TCRB and IGH in patients is significantly shorter than that in healthy controls, which is the result of both VDJ rearrangement and clone selection. We also found that the IgA1 frequency in the PBMC of IgAN is significant higher than that in other Nephropathy (NIgAN) and healthy control, which is consistent with the previous reports on the level of IgA1 producing B cells and serum IgA1. Significantly, we identified a set of IgAN disease related TCRB and IGH CDR3s, which can be used to distinguish IgAN from NIgAN and healthy controls from the blood with high accuracy. These results indicated that TCRB and IGH repertoire can potentially serve as non-invasive biomarkers for IgAN diagnosis. The characteristics of kidney infiltrating and circulating lymphocytes repertoire shed light on IgAN detection, treatment and surveillance.

## Introduction

Immunoglobulin A Nephropathy (IgAN) is the most common glomerulonephritis, especially in South East Asia (1)(2). In Singapore, it accounts for about 40% of the primary glomerulonephritis (2); in China, IgAN accounts for about 40-50% of the primary glomerular disease (3)(4)(5). Only 5-30% of the IgAN patients could get a complete remission, and about 30-50% patients will develop an end-stage renal disease within 20 year of the disease initiation (1), which damages the glomeruli badly and brings in a considerable physical agony and economic burden to the patients and the families.

The disease derived its name from the deposition of immunoglobulin A (IgA), especially IgA1 subclass, in the kidney, which distinguishes IgAN from other glomerulonephritis without IgA deposits (NIgAN). However, the exact pathogenesis of IgAN is still incompletely understood. Lai (6) and Suzuki *et al*.(7) reviewed the previous studies and discussed the pathogenesis of IgAN in details, and they considered that four hits are needed for the kidney injure of IgAN patients. The first hit is the production of galactose-deficient IgA1 (8)(9)(10), which has been reported widely to be genetically determined (11)(12); the second hit is the production of antiglycan antibodies, which mainly is the IgG and IgA subtypes, and can specifically bind the under galactosylated IgA1; the third hit is formation of IgA immune complex, which then deposit in the mesangium; the fourth hit is the activation of mesangial cells by immune complex, which can proliferate and secrete extracellular matrix, cytokines, and chemokines, and then activate the alternative complement pathway and result in renal injury. In fact, it is widely accepted that the IgAN is an immune related disease with the involvement of the whole immune system, and except of the B cells and antibody, the T cells also play important roles in the disease pathogenesis.

The definite diagnosis of IgAN relies on the histopathology of renal biopsy (13), which showed IgA deposition on the glomerular mesangium according to immunofluorescence and electron microscopy. However, renal biopsy is invasive and could bring in some possible risk, such as developing an infection and potential damage to the kidney, and it is impractical to perform renal biopsy repeatedly during and after treatment to monitor the disease status and prognosis. Therefore, there is an urgent need for reliable noninvasive biomarkers to diagnose subclinical IgAN, to estimate the disease activity and to assess the treatment efficacy (14)(15). It is also urgent to develop new therapies targeting the key pathways involved in the pathogenesis of the disease (16). Immune repertoire high throughput sequencing (HTS) has become an emerging technology to quantitatively analyze the distribution of large number of T/B cell in a biological sample (17)(18) (19). Immune repertoire HTS has been used in the fundamental research and application research of all kinds of immune related disease, such as infection disease (20), autoimmune disease (21)(22), tumor (23) and allergy (24). Disease specific T/B cells has also been used as non-invasive biomarkers in disease diagnosis and prognosis, such as tumor immunotherapy (25), leukemia minimal residual disease monitoring (26)(27), status of cytomegalovirus infection (28). Investigating the diversity of T/B lymphocytes by sequencing the receptor of those T/B cells and identifying the phenotype/antigen associated T/B cells in IgAN is critical to understand the disease and to develop novel diagnostic methods and therapies for long-term disease suppression.

In this manuscript, we analyzed the T cell receptor bata chain (TCRB) and immunoglobulin heavy chain (IGH) repertoire of kidney infiltrating and circulating lymphocytes of nephropathy patients by immune repertoire HTS, to investigate the antigen receptor characterization of IgAN comparing with NIgAN and health control. We compared every aspect including the diversity and CDR3 length of the repertoire, as well as the IGH isotype distribution and their somatic hypermutation rate between different groups. Intriguingly, we explored the possibilities to classify patients with IgAN from NIgAN and healthy controls based on the disease associated clones in PBMC.

## Methods

### Samples

This study was approved by the medical ethics committee of Xijing hospital and all specimen donors signed the written informed consent. In total, 52 nephropathy patients, including 39 IgAN and 13 NIgAN, were recruited. Clinical characteristics of these patients are showed in Table S1, and the NIgAN included membranous nephropathy, capillary disease, IgM nephropathy, glomerulosclerosis and hyperplasia glomerulonephritis. The 24 hour urine was collected in order to estimate the amount of protein excretion, and the renal function was evaluated by Urea Nitrogen, Creatinine and eGFR level. Lee’s pathological classification was used to assess the state of the IgAN patients. The kidney tissue biopsies and peripheral blood (PB) were collected during the disease diagnosis. We also recruited 60 healthy volunteers without history of cancer, autoimmune disorder and surgery, and 5ml PB were drawn from each healthy volunteer. All biopsies were frozen in liquid nitrogen immediately and were stored in -80 °C until usage. For PB samples, the peripheral blood mononuclear cell (PBMC) was isolated immediately after the blood drawing using Ficoll-Paque and were also frozen in liquid nitrogen and stored in -80 °C.

### High throughput sequencing of IGH and TCRB repertoire

Total RNA was extracted from the kidney tissue and PBMC using TRIzol™ Reagent (Invitrogen, 15596026), following the manufacturer’s guidelines. We used Agilent RNA 6000 Pico Kit (Agilent, 5067-1513) to assess the quality and quantity of the total RNA. RNA of all 52 nephropathy patients and randomly selected 42 healthy controls were processed for IGH repertoire HTS. Seventeen IgAN patients with enough remaining RNA were processed for TCRB repertoire HTS, together with 18 healthy individuals used for comparison. For all samples, total RNA was used for cDNA synthesis with oligo(dT) primer using SuperScript™ II Reverse Transcriptase (Invitrogen, 18064014). In order to enrich the completely arranged IGH sequences, the single-strand cDNA reverse transcription products from 200 ng total RNA was used as templates for multiplex PCR with forward primer mix reported in our previous published paper (27) annealed to all functional variable (V) genes and reverse primer mix annealed to constant region Cα (IgA), Cμ (IgM), Cγ (IgG), Cδ (IgD) and Cε (IgE). In order to amplify the completely arranged TCRB fragments, the cDNA products from 200 ng total RNA was used as templates for multiplex PCR with forward primer mix specific for all functional V genes and reverse primer mix specific to all functional junction (J) genes. The PCR reaction conditions was the same as our previous paper (27)(29). The target amplified production (150-300 for IGH and 100-200 for TCRB) was purified by 2% agarose gel electrophoresis. The Illumina Hiseq sequence adaptors were ligated to construct sequencing libraries, which then were sequenced on Illumina Hiseq2500 platform.

### Bioinformatics analysis of the immune repertoire HTS data

Sequencing data were analyzed using the TCR and BCR repertoire analyzing pipeline *IMonitor* (30). For comparing the diversity of different samples, we randomly chose a subset of 2.5 million mapped reads for all the continuous analysis. We used the CDR3s with correct open reading frame throughout this paper unless special statement. If two samples were sequenced in the same sequencing lane, it is possible some low-frequency sequences of one sample were the polluted data from other samples due to errors of index synthesis and sequencing. Therefore, it is necessary to remove such polluted sequences especially for the disease-associated-clones analysis, and we added two steps to filtrate those polluted sequences before further analysis. Firstly, if two sequences have the same CDR3 nucleotides and above 98% similarity for the non-CDR3 part, and meanwhile the abundance ratio of this sequence between the two samples was larger than 2000:1, the sequence with lower abundance was removed from the sample. Secondly, we count the number of samples contain a certain CDR3 in a sequencing lane, and calculate the percentage of this number in the total samples number sequenced in that lane. If this percentage in one sequencing lane is twice or more than the average percentage of other sequencing lanes, the CDR3s was removed from the sample with the lowest frequency until it is nearly equal to the average percentage of other lanes.

### Classification of the IgAN patients according to the disease associated clones

#### Defining disease associated clones

In this study, we define a clone or clonotype as a unique TCRB or IGH CDR3 nucleotide sequence. Disease associated CDR3 clones were defined as those TCRB or IGH CDR3s present in at least four renal tissues of IgAN patients and in at most three healthy blood sample, in combination with the 10 most abundant CDR3s in each renal tissue of the patients. For TCRB, we also demanded the frequency of disease associated CDR3s higher than 0.1% in patients.

#### Training classification model and leave-one-out cross validation

We used the logistic regression model in R package based on two features, the proportion of unique disease associated clones presented in the sample and the proportion of total disease associated clones presented in the sample (Figure S4). Exhausted leave-one-out cross validation was used to assess the identifier’s performance during model training. Concretely, given there are N samples with IgAN, N-1 samples were used as training data and process the above classification model. The left one sample was used as testing data to perform the classification. The cross validations were repeated for 5N times until every sample was used as testing data for 5 times.

### Statistical analysis

For comparison of the two groups, the Mann-Whitney U-test was used unless otherwise noted. *, ** and *** indicate P values less than 0.05, 0.01 and 0.001 respectively. In box and whisker plots, the box extends from the 25th to 75th percentiles, the line in the middle represents the median and outliers represent the minimum and maximum value.

## Result

### Clinical and pathological characteristics of the patients

The clinicopathological data of the patients is summarized in table 1. For IgAN patients, 30 of them are male, and 9 are female. As to NIgAN patients, 8 are male and 5 are female (Table 1). Most of the IgAN patients were diagnosed as grade of 2, 3 or 4 according to the Lee’s classification. The average 24-hours proteinuria was 2169 mg for patients with IgAN, and varied largely among patients, with 5 patients have a urine protein above 5 g/day. The urea nitrogen, creatinine and eGFR also varied extensively among patients, with an average level of 6.17 mmol/L, 111.51 μmol/l and 81.56 ml/min/1.73m2 respectively (Table S1). We did not observe any positive correlations between those renal function related clinical indices with the Lee’ classification (Table 1). When we compared IgAN with NIgAN, no significant differences were observed for those clinical indices (Figure S1), which reflect the similarity of renal function between patients with IgAN and NIgAN.

**Table 1.** Summary of clinical and laboratory information of the recruited patients.

| Label | Lee' C | Sex<br>(M vs F) | Age | Proteinuria<br>(mg/24h) | S. Cre.<br>($\mu\text{mol/l}$) | S. Urea N.<br>(mmol/l) | eGFR<br>(ml/min/1.73 m<sup>2</sup>) |
| --- | --- | --- | --- | --- | --- | --- | --- |
| Average (range) |  |  |  |  |  |  |  |
| IgAN | 1 | 2 vs 1 | 23.7<br>(15-29) | 5824.7<br>(3364-7810) | 110.7<br>(58-152) | 8.9<br>(4.0-12.9) | 82.3<br>(58-119) |
| IgAN | 2 | 10 vs 2 | 29.8<br>(15-68) | 1849.4<br>(185-8250) | 101.7<br>(76-168) | 5.3<br>(3.3-10.4) | 89.9 (60.4-130.2) |
| IgAN | 3 | 8 vs 4 | 29.8<br>(9-52) | 2016.5<br>(700-4030) | 108.2<br>(54-212) | 6.6<br>(3.0-10.7) | 81.1 (37.3-145.6) |
| IgAN | 4 | 8 vs 2 | 35.2<br>(21-51) | 2124.9<br>(157-6570) | 126.3<br>(64-295) | 6.0<br>(3.39-8.7) | 73.0<br>(22.7-132) |
| IgAN | 5 | 2 vs 0 | 35.5<br>(33-38) | 1568.5<br>(97-3040) | 112<br>(97-127) | 5.2<br>(4.2-6.3) | 80.1<br>(63.5-96.7) |
| NIgAN | NA | 8 vs 5 | 36.8<br>(14-57) | 1698<br>(95-4510) | 90.0<br>(62-132) | 5.1<br>(3.3-7.3) | 86.6 (59.6-127.3) |

Lee's C, Lee's classification; 24 h Prot., S. Cre., Serum Creatinine; S. Urea N., Serum Urea Nitrogen.

### IGH and TCRB in the renal tissue of the patients with IgAN showed decreased repertoire diversity compared with that in PBMC

VDJ recombination and other factors introduced an astronomical diversity in the CDR3 sequences. Therefore, the CDR3 sequence could serve as a natural barcode for a particular IGH or TCRB clone. We measured the IGH and TCRB diversity by three indices, the number of unique CDR3, the Shannon index and the Gini coefficient of CDR3. Both the IGH and TCRB in renal tissue of patients with IgAN were less diverse than that in PBMC of patients with IgAN and healthy controls. Concretely, the number of unique CDR3 was approximately fivefold less in renal tissue than that in PBMC (Figure 1A, D); the Shannon index of CDR3s in renal tissue was significant lower than that in PBMC (Figure 1B, E); the Gini coefficient of CDR3s was significant higher in renal tissue than that in PBMC (Figure 1 C, F). All these three indices indicated that the IGH and TCRB repertoire in the renal tissue was significantly less diverse than that in PBMC, which resulted from both the clonotypes reduction and their imbalanced frequencies in renal tissue compared with PBMC. When comparing the TCRB and IGH diversity of PBMC between patients with IgAN and healthy controls, no significant difference was observed (Figure 1). We also performed IGH repertoire analysis for 13 patients with NIgAN; however, no significant difference for IGH diversity was observed between IgAN and NIgAN in both renal tissue and PBMC (Figure 1 D, E, F). We then tried to investigate if the TCRB and IGH repertoire diversity could be related with some clinical indices, including 24-hours proteinuria, serum creatinine, urea nitrogen, eGFR and Lee’s grade, but no strong correlations were observed (Figure S2A, S3A).

**Figure 1.**
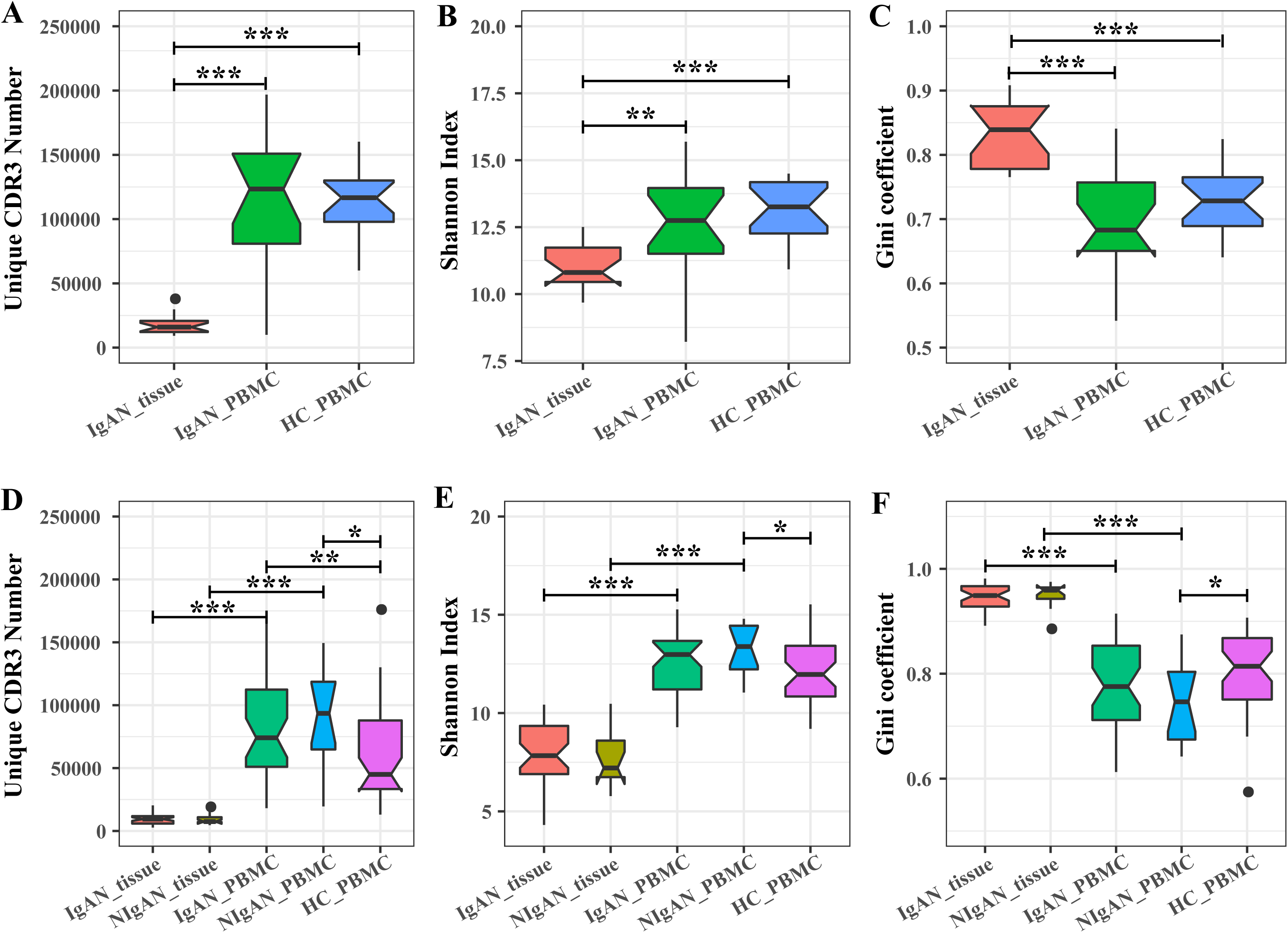
The TCRB (A-C) and IGH (B-D) CDR3 diversity in renal tissue and PBMC of patients and healthy controls. The diversity was measured by the unique CDR3 number (A, D), Shannon index (B, E) and Gini coefficient (C, F). HC, healthy controls.

### Increased IgA1 frequency and decreased IgG frequency in PBMC for patients with IgAN

We compared the distribution of IgA, IgG and IgM isotypes between tissue and PB, and between patients and healthy controls. For IgA, we subdivided it into IgA1 and IgA2 to see the percentage of the two different IgA transcripts. The frequency of IgA1 in PBMC was significant greater for IgAN patients than NIgAN patients and healthy control (Figure 2A). Although more IgAN patients have higher IgA1 percentage in the renal tissue than NIgAN patients, no significant difference was observed (Figure 2A). For IgG, in contrast to IgA1, its frequency in PBMC was significantly lower for IgAN patients than for NIgAN patients and healthy control, but there is no significant difference for its frequency in renal tissue between patients with IgAN and NIgAN (Figure 2B). For IgA2 and IgM, no significant difference was observed between any two groups (Figure 2C, 2D). The IgA1 and IgG frequencies in the renal tissue were significantly higher than that in PBMC for both IgAN and NIgAN patients (Figure 2A, C). No obvious correlations were observed between the IgA1 frequency in PBMC with the clinical indices and lee’s grade (Figure S3B).

**Figure 2.**
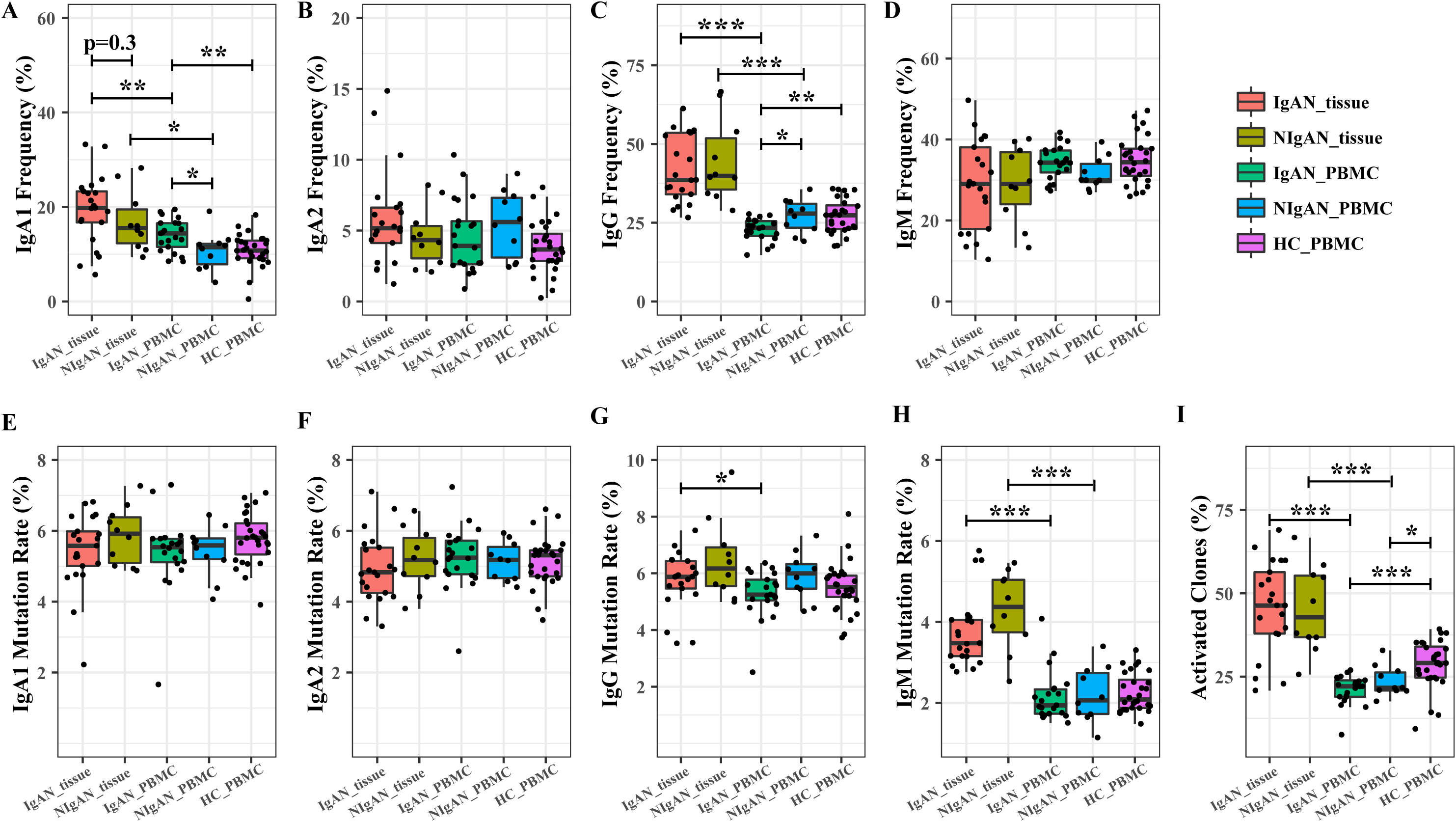
The isotype distribution and somatic hypermutation of each IGH isotype in renal tissue and PBMC of patients and healthy controls. (A-D) IgA1 (A), IgA2 (B), IgG (C) and IgM (D) frequency distribution in renal tissue and PBMC of patients and healthy controls. (E-H) IgA1 (E), IgA2 (F), IgG (G) and IgM (H) somatic hypermutation rate in renal tissue and PBMC of patients and healthy controls. The activated clone frequency (I) in renal tissue and PBMC of patients and healthy controls. HC, healthy controls. The P value of IgAN_tissue vs IgAN_PBMC, NIgAN_tissue vs NIgAN_PBMC, IgAN_tissue vs NIgAN_tissue, IgAN_PBMC vs NIgAN_PBMC, IgAN_PBMC vs HC_PBMC, and NIgAN_PBMC vs HC_PBMC was calculated.

### B cell activation in the renal tissue of disease

Class switching and somatic hypermutation (SHM) are two biological mechanisms through which B cells adapt to pathogens. During B cell activating and responding to antigens, the naïve IgM B cells switch to IgA, IgG or IgE B cells, and accumulate SHMs. To investigate the B cell activation in IgAN, we defined activated IGH clones as those IgA or IgG with high mutation rate (above 3%) as reported previously (31). The SHM rate of each IGH clone was calculated by determining the average mismatch rate per base-pair unit in the non-CDR3 region. SHM rate of IgA did not show significant difference between any two groups (Figure 3E, F). IgG in kidney tissue have higher SHM rate than that in PBMC for patients with IgAN, however there is no significant difference between the tissue and PBMC for patients with NIgAN (Figure 2G). IgM from renal tissue of patients displayed a higher average SHM rate than IgM from PBMC (Figure 2H). When comparing the SHM rate between the PBMC of patients and healthy controls, no significant difference was observed for any isotype. We then investigate the activated clone frequency in renal tissue and PBMC of patients, as well as in healthy controls. The result demonstrated that the frequency of activated IGH clones was significant higher for B cells infiltrating in the renal tissue than in PBMC (Figure 2I). We also found that the activated IGH clones proportion in the PBMC of both IgAN and NIgAN was significant lower than that in the PBMC of healthy control. We did not observe obvious difference for the frequency of activated IGH clones between IgAN patients and NIgAN patients in both the renal tissue and PBMC (Figure 2J).

**Figure 3.**
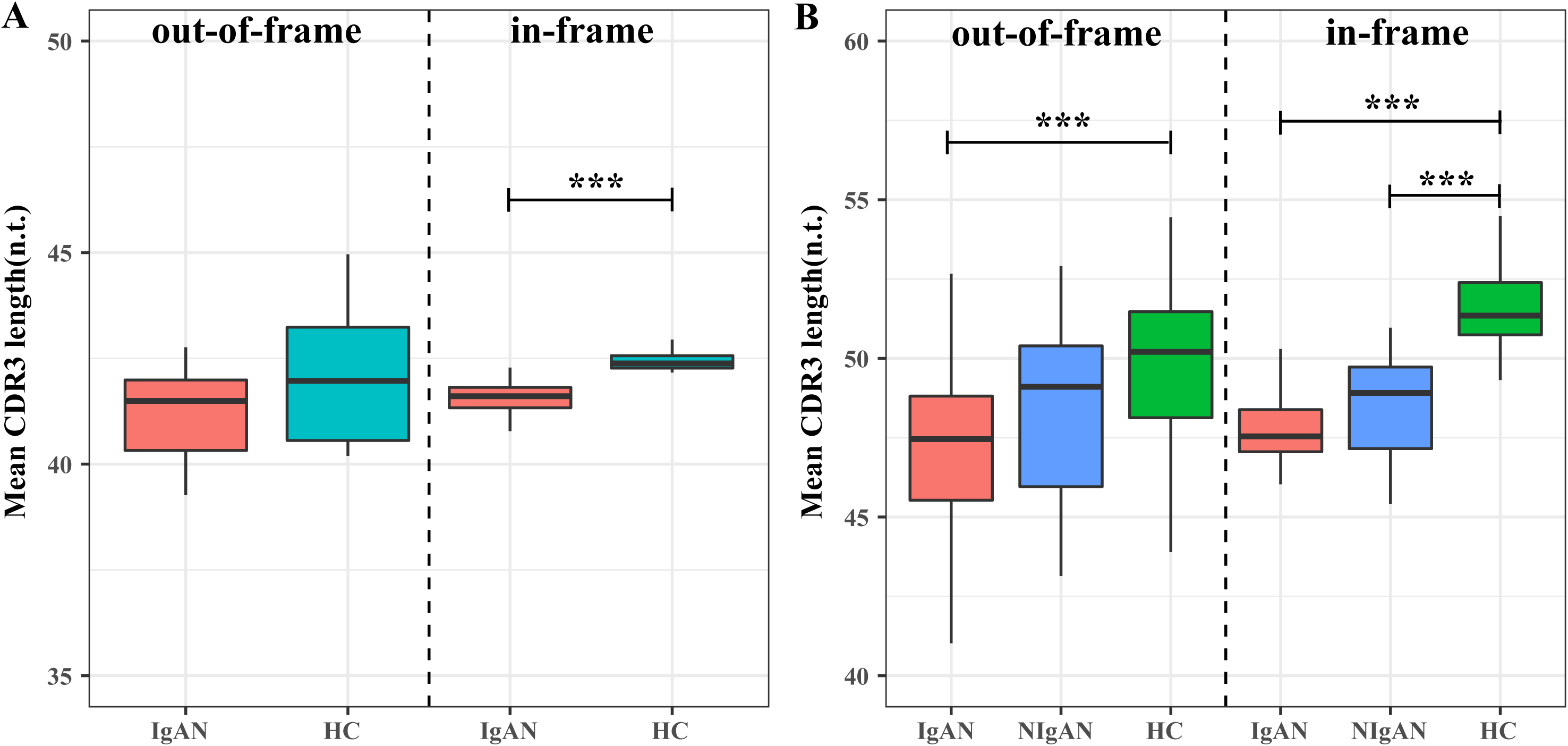
TCRB and IGH CDR3s length distribution in IgAN patients and healthy controls. The mean CDR3 length of out-of-frame (left panel) and in-frame (right panel) TCRB (A) and IGH (B) in the PBMC of patients and healthy control. HC, healthy control. n.t., nucleotide.

### TCRB and IGH reveal shorter CDR3 length in IgAN

We observed a significant reduction in both TCRB and IGH CDR3s nucleotide length in PBMC of IgAN patients comparing with healthy controls (Figure 3A, B, right panel). The shorter TCRB and IGH CDR3s could either be a feature of the initial VDJ rearrangement, or the result of negative and positive selection during T/B cell development/maturation and antigenic selection in periphery, or the combination of both. To investigate the origination of the shorter CDR3s in patients, we analyzed the CDR3 length of the out of frame TCRB and IGH sequences, which did not go through the negative and positive selection, as described by previous studies(32)(22)(33). The out-of-frame TCRB CDR3s in IgAN patients and heathy control had similar length, (Figure 3A, left panel), indicating the shorter TCRB CDR3s in IgAN patients is mainly the result of the positive and negative selection during T cell development in the thymus. For IGH, the pre-selected out-of-frame CDR3s in IgAN patients were also shorter than that in healthy controls (Figure 3B, left panel), however, the in-frame functional clones showed more marked distinction between IgAN patients and healthy controls (Figure 3B, right panel). Therefore, the shorter IGH CDR3 in patients is the combined result of increased production of clones with shorter CDR3 during VDJ rearrangement and the clone selection during lymphocytes maturation and in periphery.

### Disease associated clonotypes

We hypothesis that a set of pathogenic or disease associated T and B cells exist in the renal tissue, and are particularly interested in their appearance in the blood and the potentiality to serve as non invasive biomarkers for IgAN diagnosis. Firstly, we evaluated the repertoire overlap between the renal tissue and the blood. The overlap rate was calculated by adding together the total number of CDR3s identified in the both repertoires, and dividing it by the sum of CDR3 number of the two repertoires. The overlap rate between the renal tissue and PBMC repertoire of the same patients was significantly higher than the inter-individual rate, for both TCRB and IGH (Figure S4A, C). Additionally, an average of 34.13% of TCRB CDR3s and 30.21% of IGH CDR3 from the renal tissue can be found in the PBMC of the same patient (Figure S4B, D). In several samples, more than 60% of renal tissue TCRB and IGH CDR3 can detected in PBMC. The frequent appearance of renal tissue lymphocytes in PBMC suggested the diagnostic potential of the blood. Considering the antigenic complexity and the background noise in the blood, we intended to search the disease associated CDR3 clonotypes in the renal tissue, and further used those in the blood to do the diagnosis. Disease associated CDR3s are usually thought to be widely and abundantly represented in the pathogenic tissues but scarcely appear in the healthy tissues (28). Accordingly, IgAN associated T and B cell clonotypes were selected as described in the methods. We identified 312 TCRB CDR3s and 462 IGH CDR3s suggested to be associated with IgAN (Table S2 and S3). Further evidences supporting the associations were identified. Firstly, the abundance of the associated CDR3s exclusive of the 10 most abundant ones in each renal tissue, is significant higher than the other CDR3s (Figure 4A, 4C), which suggested those clones had been activated and undergone expansion. Secondly, we found that the disease associated TCRB and IGH CDR3s were significant shorter than the other CDR3s (Figure 4B, 4D), which is in accordance with the reported feature of autoimmune T and B cells (22)(34). Interestingly, we found that the unique frequencies (the number of unique disease associated CDR3s divided by the number of total unique CDR3) of disease associated TCRB and IGH CDR3s were positively correlated, in both the renal tissue and the blood (Figure S5A, B), which may due to the specificity of TCRB and IGH CDR3 when recognizing IgAN antigens. We also tried to investigate if the total frequency of those public clones is related to the clinical indices and lee’s grade, however no obvious correlation was observed (Figure S2B, S3C).

**Figure 4.**
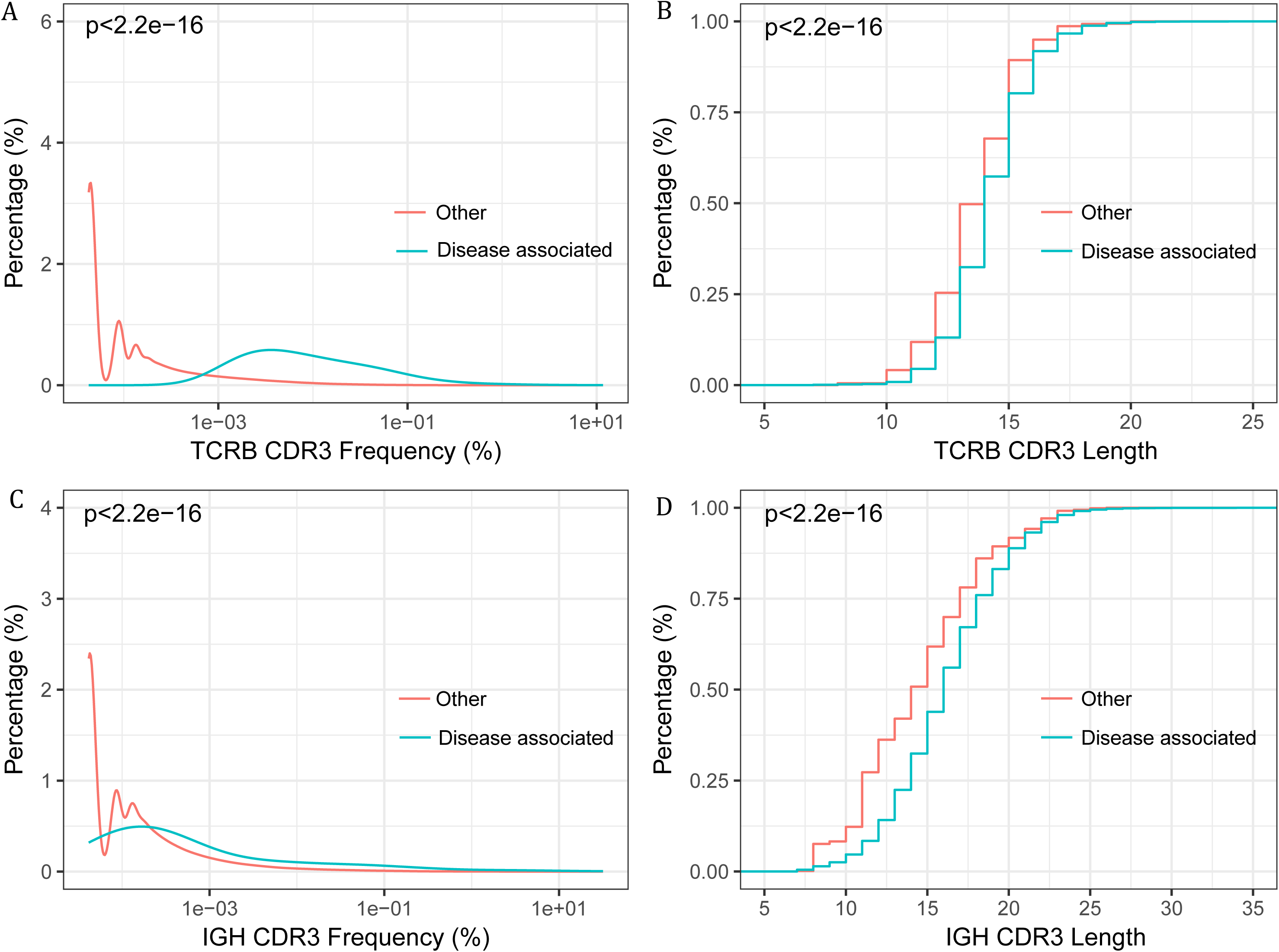
The frequency (abundance) and CDR3 length of the disease associated CDR3 clones and other clones. (A) The frequency distribution of disease associated TCRB CDR3 versus other TCRB CDR3s. (B) Cumulative length plots of the disease associated TCRB CDR3 versus other TCRB CDR3s. (C) The frequency distribution of the disease associated IGH CDR3 versus other IGH CDR3s (D) Cumulative length plots of the disease associated IGH CDR3 versus other IGH CDR3s.

### Noninvasive classification of IgAN patients with the disease associated lymphocytes

Next, we intended to use the identified IgAN associated CDR3 to do the noninvasive classification of IgAN from the PBMC data. Firstly, we evaluated and found that both the unique and total disease associated TCRB CDR3 frequencies in PBMC were significantly higher in patients with IgAN than in healthy controls (Figure 5B, C); for disease associated IGH CDR3s, there were also obvious differences for their unique frequency and total frequency between IgAN patients and healthy controls, and between IgAN patients and NIgAN patients (Figure 5D, E). Then we constructed a binary classifier to distinguish the IgAN patients from healthy controls utilizing the unique and total frequency of disease associated TCRB or IGH CDR3s as features. The disease associated TCRB CDR3 can separate the PBMC of IgAN patients and healthy control with an accuracy of 94.11%, (AUC (the area under the ROC curve) = 96.08%) (Figure 5F), which indicated that those disease associated TCRB clones could be used as potential biomarkers for IgAN. In contrast, the disease associated IGH CDR3 supported a worse performance with an accuracy 74.87% (AUC = 75.61%) (Figure 5G). We also tried to see if those disease associated IGH clones can be used to distinguish the IgAN patients from NIgAN patients, the result showed the accuracy of discrimination was 78.46% (Figure 5H), which indicated that those IgAN associated IGH clones were more prone to exist mainly in patients with IgAN than NIgAN. Overall, we demonstrated that IgAN could be well classified noninvasively by TCRB and IGH repertoire data.

**Figure 5.**
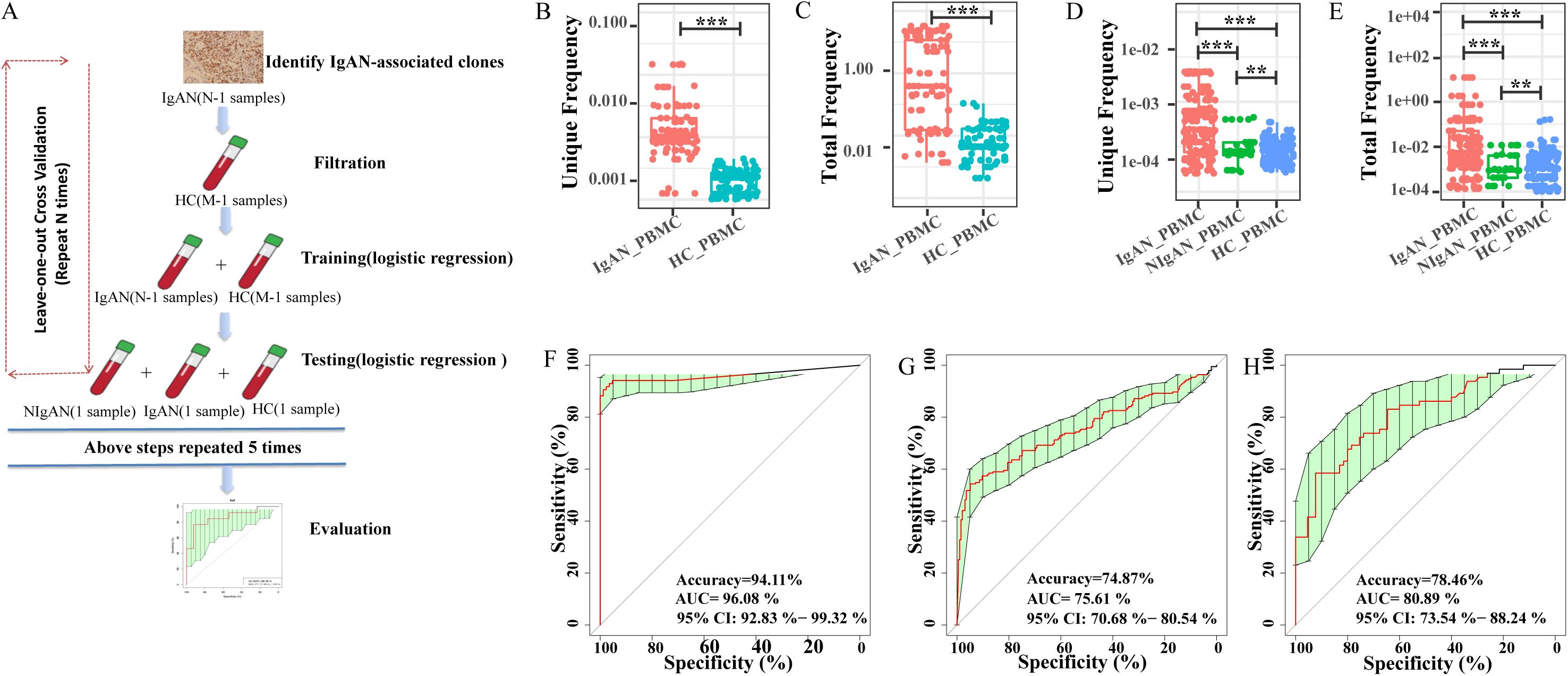
Classifying different groups according to TCRB and IGH repertoire data. (A) Analytical overview, which was described in detail in the methods. (B-E) The burden of disease associated clones in PBMC of different groups. The unique frequency (B) and total frequency (C) of disease associated TCRB CDR3s in PBMC of IgAN patients and healthy controls. The unique frequency (D) and total frequency (E) of disease associated IGH CDR3s in PBMC of IgAN patients, NIgAN patients and healthy controls. (F-H) The leave-out-one cross validation ROC curves of classification according to the disease associated clones. The ROC curve of classification of patients with IgAN and healthy controls according to disease associated TCRB CDR3s (F) and disease associated IGH CDR3s (G). The ROC curve of classification of patients with IgAN and NIgAN according to disease associated IGH CDR3s (H). HC, healthy controls.

## Discussion

In this study, we performed TCRB and IGH repertoire HTS of paired renal tissue and PBMC of patients with IgAN and NIgAN, as well as of PBMC of healthy controls, to investigate the pathogenesis of IgAN and develop new biomarkers for the disease. As far as we know, this is the first study focusing on both the T and B cell receptor repertoire of IgAN by HTS. Comparing with the lymphocytes in peripheral blood, renal tissue infiltrating lymphocytes was less diverse (Figure 1), which indicated that kidney infiltrating lymphocytes contained more clones undergone clonal expansion due to antigen stimulation. The clonal expansion of γδ T cells in the kidney tissues comparing with PBMC has been reported by others (32). In our study, higher frequency of IgA and IgG, and the increased IgM SHM rate comparing with PBMC (Figure 2) also supports the universal activation of the tissue infiltrating lymphocytes.

A recent study indicate aberration in TCR VDJ recombination confer risks to autoimmune diseases such as type 1 diabetes, in which the diabetes autoantigen specific T cells were found to have shorter TCRB CDR3 than viral specific T cells and global T cells (21). We also observed that the average lengths of both TCRB and IGH CDR3 in IgAN patients were shorter than those in healthy controls, from the PBMC data (Figure 3). By investigating separately the out-of-frame and the in-frame CDR3s, we conclude that VDJ recombination and selection during T/B cell development and in periphery play different roles in TCRB and IGH. Additionally, the IgAN associated TCRB and IGH clonotypes also showed significantly shorter CDR3 length. Our finding provide further evidence that the generation of a self-reactive prone T and B cell repertoire is account for the susceptibility of common autoimmune diseases.

Human IgA has two subclasses, IgA1 and IgA2, and IgA1 is a functionally and structurally distinct immunoglobulin from IgA2 (33). Numerous studies have demonstrated that the IgA deposited in the glomeruli of patients with IgAN is the subclass IgA1(34)(35)(6), which includes three to six O-linked glycan chains in the hinge region between the first and second constant region. One possible reason for the increased IgA1 immune complex is the overproduction of IgA1, which could be assessed from the level of IgA1 producing B cell, the IgA1 mRNA and the IgA1 antibody in the serum. Several studies have reported that the percentage of IgA1 producing B cell was higher in PB of patients with IgAN than health controls (36)(34). We found that the IgA1 frequency in PBMC of patients with IgAN was significant higher than patients with NIgAN and healthy controls (Figure 2), which may due to more IgA1 producing B cells or higher transcription activity of those B cells. A number of literatures have reported a higher serum IgA level in IgAN (37)(38), and the association of the serum galactose-deficient IgA1 level with disease progression (39). The above data indicate that the IgA1 level could be used as clinical biomarkers to assist the diagnosis and prognosis of IgAN.

Although the serum level of IgA1 elevates in most of the patients with IgAN, the accuracy of using serum IgA1 or galactose-deficient IgA1 as diagnostic standard to replace the histological examination of renal biopsy is insufficient. In fact, due to lack of single or combined laboratory test, screening for IgAN is not feasible yet. Therefore, searching for new biomarkers in liquid biopsies, such as blood or urine, is urgently needed. Because IgAN is an immune related disease, there should be abundant disease associated T/B cells targeting the disease related autoantigens infiltrating in the injured tissues, and our data (Figure S4B, D) and published paper (34) suggested those disease associated T/B cells derive from the bone marrow and circulate in the PB. In this paper, we identified a set of disease associated TCRB and IGH CDR3 from the renal tissues which then was used to distinguish the peripheral blood of IgAN patients from healthy individuals. With the disease associated IGH CDR3 panel, the leave-out-one cross validation showed an accuracy of 74.87% when classifying IgAN patients and healthy individuals (Figure 5G); more importantly, this IgAN associated IGH CDR3 panel can also distinguish patients with IgAN from NIgAN with an accuracy of 78.46% (Figure 5H). What is striking is that the disease associated TCRB CDR3 panel can predict the IgAN status from healthy controls with a very high accuracy of 94.11% (Figure 5F). In fact, some studies have reported the involvement of T cell in the pathogenesis of IgAN, such as increased circulating αβ and γδ T cells in PBMC of IgAN patients (40)(41), involvement of T cell in the progression of IgAN (42), and the existence of conserved TCRB CDR3 in the IgAN biopsies (43). Overall, our study further demonstrates that disease associated TCRB and IGH CDR3 could work as the potential biomarker to help the diagnosis of IgAN, at least in screening the IgAN. However, due to the limited samples in our study, validating the performance of those biomarkers and demonstrating the accuracy and reproducibility of this method by more studies from other laboratories is needed.

In summary, we present here a comprehensive landscape of T and B cell repertoire in IgAN. We reported their interesting features, including the restricted repertoire size and increased activation of kidney infiltrating lymphocytes, the increment of IgA1 frequency, and shorter CDR3 length for both TCRB and IGH of IgAN patients, which may relate with the autoimmune nature of those lymphocytes. The most important discovery is that using our defined disease associated TCRB and IGH CDR3s, we can distinguish PBMC of IgAN patients from healthy controls. These findings demonstrate the potentiality of disease related T/B cells as alternative biomarkers for the screening, diagnosis and prognosis of IgAN.

## Acknowledgement

This work was supported by the National Natural Science Foundation of China (No. 81672593 and No. 81670655), the Major Nature Science Foundation of Shaanxis Province (No. 2017ZDXM-SF-045), and Shenzhen Municipal Government of China (JCYJ20170817145404433).

---

**Figure S1.** The comparison of clinical data between IgAN and NIgAN patients. The age (A), 24-hours proteinuria (B), serum creatinine (C), serum urea nitrogen (D) eGFR of patients. eGFR, estimated glomerular filtration rate.

**Figure S2:** The association of clinical indices with TCRB CDR3 Shannon index (A) and total disease associated TCRB CDR3 frequency (B) in the PBMC of patients.

**Figure S3:** The association of clinical indices with IGH CDR3 Shannon index (A), IgA1 frequency (B) and total public TCRB CDR3 frequency (C) in the PBMC of patients. Red dot, IgAN; blue dot, NIgAN.

**Figure S4.** CDR3 Overlap rate of two repertoires. TCRB CDR3 (A) and IGH CDR3 (C) Overlap rate of any two samples of different patients and overlap rate of renal tissue and PBMC of the same patients. The percentage of overlap TCRB CDR3 (B) and IGH CDR3 (D) between renal tissue and PBMC of the same patients in renal the tissue.

**Figure S5.** The correlation of the unique frequency between disease associated TCRB and IGH CDR3 in renal tissue (A) and the PBMC (B).

